# Integration of heterogeneous functional genomics data in gerontology research identifies genes and pathway underlying aging across species

**DOI:** 10.1101/451013

**Authors:** Jason A. Bubier, George L. Sutphin, Timothy J. Reynolds, Ron Korstanje, Axis Fuksman-Kumpa, Erich J. Baker, Michael A. Langston, Elissa J. Chesler

**Author notes:** Corresponding author (EJC). Authors contributed equally to the manuscript (Jason A. Bubier, George L. Sutphin, Timothy J. Reynolds).

## Abstract

Understanding the biological mechanisms behind aging, lifespan and healthspan is becoming increasingly important as the proportion of the world's population over the age of 65 grows, along with the cost and complexity of their care. BigData oriented approaches and analysis methods for integrative functional genomics enable current and future bio-gerontologists to synthesize, distill and interpret vast, heterogeneous data. GeneWeaver is an analysis system for integration of data that allows investigators to store, search, and analyze immense amounts of data including user-submitted experimental data, data from primary publications, and data in other databases. Aging related genome-wide gene sets from primary publications were curated into this system in concert with data from other model-organism and aging-specific databases, and used in several application using GeneWeavers analysis tools. For example, we identified *Cd63* as a frequently represented gene among aging-related genome-wide results. To evaluate the role of *Cd63* in aging, we performed RNAi knockdown of the *C. elegans* ortholog, *tsp-7*, demonstrating that this manipulation is capable of extending lifespan. The tools in GeneWeaver enable aging researchers to make new discoveries into the associations between the genes, normal biological processes, and diseases that affect aging, healthspan, and lifespan.

## Introduction

The population of individuals aged 65 and over is projected to be approximately 83.7 million in 2050, almost double its estimated number of 43.1 million in 2012 [1]. Aging affects the entire organism, with age-associated decline occurring across organ systems and within distinct tissues and cell-types. One approach to identifying shared mechanisms of aging is to make systems-level comparisons at the molecular level. To discover mechanistic pathways that may have applications in prediction and extension of life-and health-span, researchers have been analyzing the biological process of aging using a variety of high-throughput technologies including genome sequencing, RNAseq, proteomics, and structural biology. These studies make use of diverse animal model systems, such as nematotodes, fruit-flies, mice and rats to characterize natural aging, and drugs or environmental interventions that affect longevity.

There are several representative applications that merit an integrative genomics approach to aging. One application is to determine which molecular and cellular factors responsible for the process of cellular senescence also underlie functional cognitive decline. Cellular senescence is an anti-cancer and wound healing mechanism characterized by arrested cellular proliferation and secretion of pro-inflammatory cytokines, chemokines, growth factors, and proteases (the senescence associated secretory phenotype, or SASP). Senescent cells accumulate with age in many tissues, where the SASP promotes chronic inflammation and exacerbates age-associated degeneration and hyperplasia. Recent evidence suggests that neurological aging and neurodegeneration are accompanied by an accumulation of secretory cells in brain, suggesting that cellular senescence may contribute to brain aging [2] through a shared mechanism. Using functional genomics studies of both the biology of cellular senescence and cognitive aging, overlapping mechanisms can be detected.

A second application is to determine what gene products are common to, two different disease states, as has been observed for obesity and dementia [3]. An integrative functional genomics approach can be used to determine the common molecular and cellular bases of these seemingly different disease conditions. A third application is to identify molecular mechanisms common to multiple anti-aging interventions. Dietary restriction has been shown to effectively increase lifespan in a variety of species [4]. An ongoing effort is underway to identify drugs that mimic the beneficial outcomes by targeting molecular pathways downstream of dietary restriction. A number of pharmacological compounds have been identified that are capable of extending the life span of invertebrates and rodents, and represent potential dietary restriction mimetics [5]. Using heterogeneous data integration, large data sets can be investigated to determine whether a common network of genes underlies both approaches to life extension, that of dietary restriction or pharmacological intervention. Finally, in a fourth application one may determine whether a gene(s) function in aging has an evolutionarily conserved role. Relying upon the assumptions of phylogenomics, whereby an ortholog of one species has the same function in ancestral species, we can use large scale data from multiple species to understand conserved mechanisms of disease. Many additional application of global gene set integration exist.

With the goal of addressing these types of questions, and in supporting the research community in applying integrative functional genomics analysis of datasets obtained across different types of experiments and organisms, we created GeneWeaver [6]. This freely available web-based software system for the collection and analysis of functional genomic experiments allows for the rapid and easy integration of large quantities of heterogeneous data. We have been curating the aging literature, and applying integrative functional genomics to identify relations among biological pathways of aging (Table 1).

**Table 1.** Summary statistics on a search of GeneSets, genes, or ontologies for “senescence OR aging OR longevity.”

| Types of GeneWeaver Data | Number of GeneSets |
| --- | --- |
| Aging-related sets | 1,655 |
| GO<sup>1</sup> term-based | 71 |
| MP<sup>2</sup> term-based | 63 |
| HPO<sup>3</sup> term-based | 109 |
| MeSH<sup>4</sup> term-based | 231 |
| QTL<sup>5</sup> | 409 |
| GWAS<sup>6</sup> | 5 |
| Gene expression | 169 |
| Drug-regulated gene sets [7] | 253 |
| Genes co-expressed to the aging phenotype | 18 |

<sup>1</sup>GO (Gene Ontology), <sup>2</sup>MP (Mammalian Phenotype Ontology), <sup>3</sup>HPO (Human Phenotype Ontology), <sup>4</sup>MeSH (Medical Subject Headings), <sup>5</sup>QTL (Quantitative Trait Loci), <sup>6</sup>GWAS (Genome Wide Association Study).

The data from these large-scale approaches provide a means to compare and contrast aging across model systems, tissues and interventions. Unfortunately, much data is presented in a non-computable format such as in primary publications, or in diverse and Balkanized data resources that require extensive efforts for integrative reanalysis. For example, the supplemental tables in publications often include lists of genes or gene networks that are analyzed in the publication but never fully integrated with existing data collected in related contexts. There have been several successful efforts to address these problems in the aging field, with large-scale, aging-related-omics databases including GenAge [7] and AgeFactDB [8]. These are excellent data repositories, however, and it remains difficult to integrate heterogeneous data across studies, organisms and experiment types. These and many other resources bring the data together in one location but lack the tools and algorithms necessary to operate on the data as a whole. Across research disciplines, but certainly also within the aging field, readily accessible data is dwarfed by the immense volume of published but uncurated data. Further, the volume of uncurated data is increasing even more rapidly. In order to harness the knowledge that lies dormant in large published datasets, two steps are necessary: 1) published data must integrated; and 2) publicly accessible tools must be developed that are capable of integrating data from multiple sources into large-scale analysis.

Here we utilize integrative functional genomics in GeneWeaver to address four questions related to aging by analyzing these large-scale, complex sets of data: a) to identify molecular relations between cellular senescence and functional cognitive decline, b) to examine the intersection between comorbid disease states, c) to identify new druggable targets for longevity, and d) to examine cross-species translation of age-related processes.

## Methods and Materials

### Integrative genomics in GeneWeaver.org

GeneWeaver is both a database of functional genomics gene sets and a suite of combinatorial and statistical tools that enable users to operate on these sets. GeneWeaver was designed to integrate large-scale genomic studies and houses these analytic tools and curated data from multiple species, ontological resources, and individual users in one central location. Users can upload sets of genes from personal or published experimental results that can then be made publicly available as part of the shared archive of gene sets in GeneWeaver, or analyzed privately. In the data archive, user submitted gene sets are integrated with gene sets from multiple sources including other individual users, publications with curated data, and large community data sources. Data resources incorporated include the drug-related gene database of the Neuroscience Information Framework (NIF) [9], GeneNetwork [10], Comparative Toxicogenomics Database (CTD) [11], Kyoto Encyclopedia of Genes and Genomes (KEGG) MeSH (Medical Subject Headings), Molecular Signatures Database (MSigDB), Online Mendelian Inheritance in Man (OMIM), and Pathway Commons. Data from these diverse resources are distilled into sets of genes within the GeneWeaver database. Additional gene sets have been created in GeneWeaver that are derived from data previously annotated to biological processes, disease states, or ontologies.

### Combine tool

The GeneWeaver “Combine” tool was used to create a set of genes representing the union of ontology annotation derived gene sets related to properties of cellular senescence (and another set of genes containing genes experimentally related to functional decline such as genome-wide differential expression data related to aging-related cognition and memory phenotypes. The combine tool was used to obtain the union of all the genes within a selected gene set and provides a count as to the number of sets each gene is found in.

### Jaccard Similarity Tool

GeneWeaver’s “Jaccard Similarity” tool was used on two genes sets to identify genes at the intersection of both senescence and functional decline. The “Jaccard Similarity” tool provides a pairwise comparison of the gene sets being analyzed. The resulting graph shows the overlap of genes in the sets using Venn diagrams. A Jaccard similarity coefficient is calculated by taking the size of the intersection of the two gene sets, and dividing by the number of unique genes available in the two sets. A value of 1.0 means perfectly overlapping, and 0 means no intersection. The “Jaccard Similarity” tool was again used to determine the similarity among mouse genes annotated to obesity and mouse genes annotated to abnormal learning and memory.

### Boolean Algebra tool

The “Boolean Algebra” tool was used to identify the genes at the intersection of two gene sets, one from each species tested for caloric restriction. A new gene set was created from the genes within this intersection. This approach allow the rapid determination of new relationships between gene sets and the creation of new gene sets based upon these findings. This same approach was repeated to compare gene sets related to differential drug treatment and overlap between both interventions.

### View Similar GeneSets

In order to identify other compounds that affect the same set of genes underlying the effects of caloric restriction, the “View Similar GeneSets” tool was used. Viewing similar gene sets within GeneWeaver compares the composition of the gene set of interest with all gene sets in the database and returns a ranked list based upon the magnitude of the Jaccard similarity of the top 250 gene sets most similar to the gene set of interest.

### GeneSet Graph

To identify the most highly connected gene within a group of gene sets related to aging, the “GeneSet Graph” tool was used. This tool presents a bipartite graph visualization of genes and gene sets. Genes are represented by elliptical nodes, and gene sets are represented by boxes. The least-connected genes are displayed on the left, followed by the gene sets, then the more-connected genes in increasing order to the right. Genes and gene sets are connected by colored lines to show what genes are in which gene sets. A degree threshold is applied on the gene partite set to reduce the graph size.

## STRING

The set of genes determined to be at the intersection of functional decline and senescence were uploaded into the STRING database (https://string-db.org/) of predicted and known protein-protein interactions. A graphical representation of the functional associations known between the encoded proteins was produced.

### Ingenuity Pathway Analysis

The genes at the intersection of obesity and dementia were uploaded to the Ingenuity Pathway Analysis software. This application maps genes to existing molecular pathways based on built in algorithms and gives predictions of for the likelihood of the observed enrichment.

### Worm Lifespan

*C. elegans* experiments were carried out using wild-type (N2) worms, originally obtained from Matt Kaeberlein at the University of Washington. The RNAi feeding clone targeting *tsp-7* was obtained from the Ahringer library [12] and the target sequence confirmed prior to use. *C. elegans* lifespan were conducted at 25°C according to standard protocols [13]. Briefly, experiments were conducted on nematode growth media (NGM) plates containing 1 mM Isopropyl β-D-1-thiogalactopyranoside (IPTG) to activate production of RNAi transcripts and 25 µg/mL carbenicillin to select RNAi plasmids and seeded with live *E. coli* (HT115) containing either *tsp-7* or *empty vector (EV)* RNAi feeding plasmids. Worms were age-synchronized via timed egg laying and transferred to plates containing 50 µM 5-fluorodeoxyuridine (FUdR) at the L4 larval stage to prevent reproduction Worms were scored as alive or dead every 1-2 days until all animals had died. Experiments were conducted in triplicate, with each replicate containing ∼105 (∼35 worms/plate on 3 plates), with each replicate showing a similar survival pattern. Survival data was pooled across replicates for statistical comparison. A Wilcoxon Rank-Sum test was applied to detect a difference in survival for worms subjected to *tsp-7(RNAi)* vs. *EV(RNAi).*

### *Cd63* in human GWAS

We examined three longevity phenotypes: mother’s age at death, father’s age at death, and the combined phenotype of parent’s age at death. GWA summary statistics were retrieved from LD Hub [14] and made available through an aging study [15] which made use of data from the UK Biobank [16]. Genomic positions for each SNP were converted to the latest genome build, hg38/GRCh38, using the UCSC liftOver tool. SNPs and positions that could not be converted were discarded. SNP reference identifiers were updated to the latest (at the time of writing) NCBI dbSNP build—version 150. SNPs without a canonical reference identifier (rsID) were discarded. Summary statistics were filtered to only include significant (p < 0.05) SNPs. Variant annotations from Ensembl v. 91 were used to annotate SNPs to the genomic features they occur in. The UK Biobank aging dataset contained five SNPs which occur in CD63. We also examined SNPs immediately downstream and upstream of CD63. There were nine downstream and 11 upstream variants. The false discovery rate, q-value was calculated using the R qvalue package [17].

## Results

### Identifying molecular relations between cellular senescence and cognitive functional decline.

GeneWeaver was queried to identify gene sets related to cellular senescence and cognitive decline. A conservative combined set of 92 genes (GS222630) unambiguously related to cellular senescence was created from the union of model organism ontology annotation to the terms MP:0008007 abnormal cellular replicative senescence, and GO:0090398 cellular senescence, GO:0090399 replicative senescence. A second set of 1,286 “experimentally observed” genes (GS222631), resulted from the complete intersection of genes from eight published studies on differentially expressed genes associated with either functional decline or aging-related cognition and memory phenotypes S1 Table [18-24]. The gene sets from each publication contain the genes the publication determined to be significantly different in each study. Using GeneWeaver’s “Jaccard Similarity” tool on these two resulting sets of genes, an intersection set of ten genes common to both senescence and functional decline was produced. Analysis in Ingenuity Pathway Analysis (IPA) reveals the gene set is enriched in MAP kinase pathway members central to many cellular pathways, including the gonadotropin-releasing hormone, neurotrophin, T-cell receptor, and B-cell receptor pathways. The ten genes were analyzed for known and predicated protein-protein interactions using the STRING database of protein-protein relations (https://string-db.org/). Five of the genes *(MapK14, Map2K1*, *Kras, Jun*, and *Hras1)* were shown to be physically interacting with be one another at the protein level. These interactions are supported by both experimental evidence and inference from the literature (Fig 1). Identification of these pathways and genes reveals that the processes of cellular senescence and functional decline share key central, conserved cellular signaling pathways in the brain.

**Fig 1.**
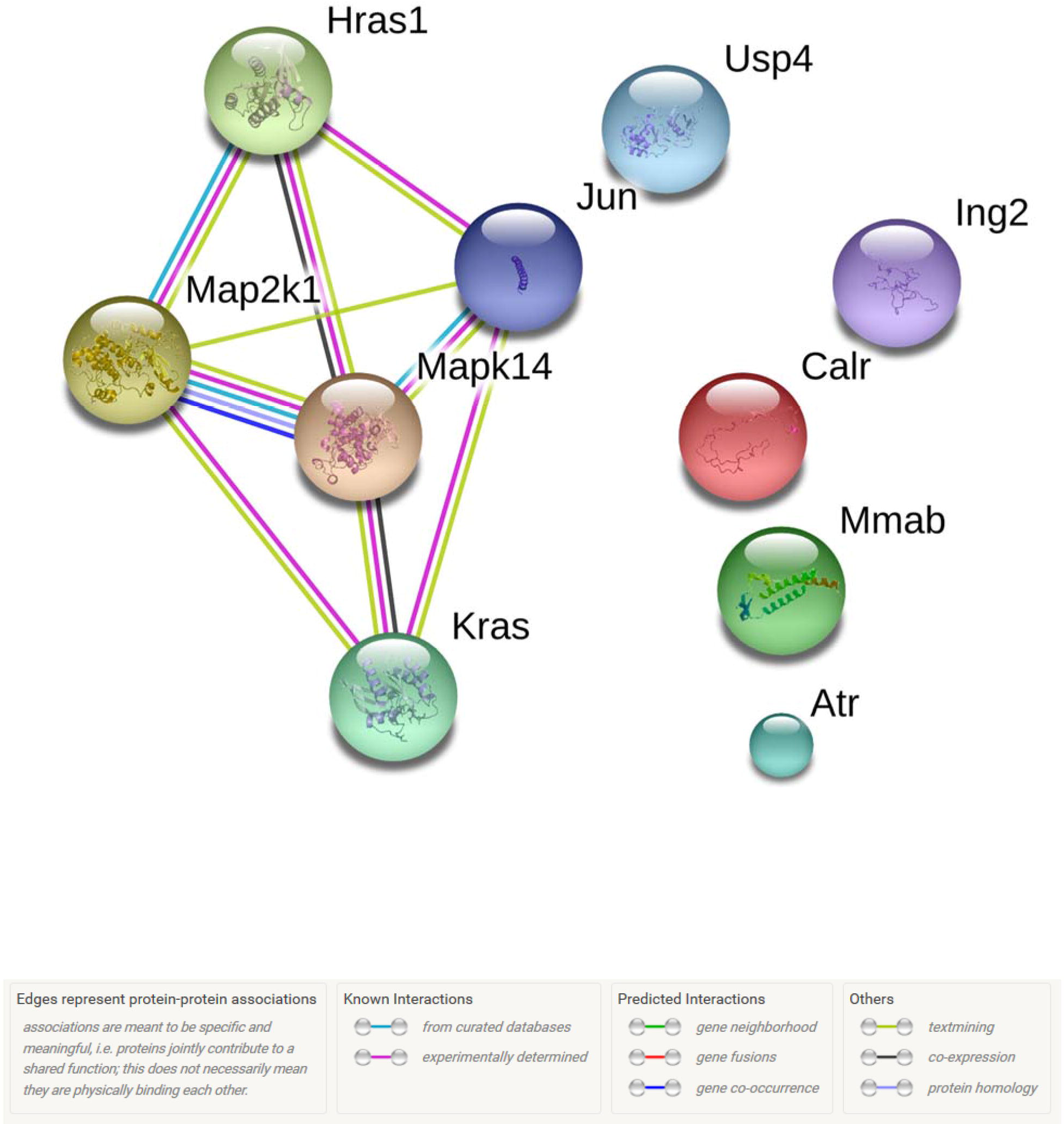
Conserved pathways between process of cellular senescence and functional decline. A group of 10 genes was identified as common to functional decline and senescence. Many of these genes interact in the MAP kinase pathway, as shown by this protein-protein interaction plot from STRING.

### Identifying a common molecular basis of obesity and dementia

To identify possible common molecular mechanisms underlying obesity and dementia, GeneWeaver database was searched to identify relevant gene sets, in this case, phenotypic alleles annotated in model organism databases to the Mouse Phenotype Ontology. Using the “Jaccard Similarity” tool in GeneWeaver, the similarity among mouse genes annotated to MP:0001261 obesity and mouse genes annotated to MP:0002063 abnormal learning and memory was determined; this overlap contained 27 genes. Ingenuity Pathway Analysis (IPA) showed that 12 of the 27 genes fall into an IPA pathway of “Behavior, Connective Tissue Development and Function, Tissue Morphology” (Fig 2). This is a highly significant enrichment (likelihood 1 x 10^31^, Fisher’s exact test). Identification of this pathway identifies potential translational targets for therapies for treating or preventing dementia in the aging obese population.

**Fig 2.**
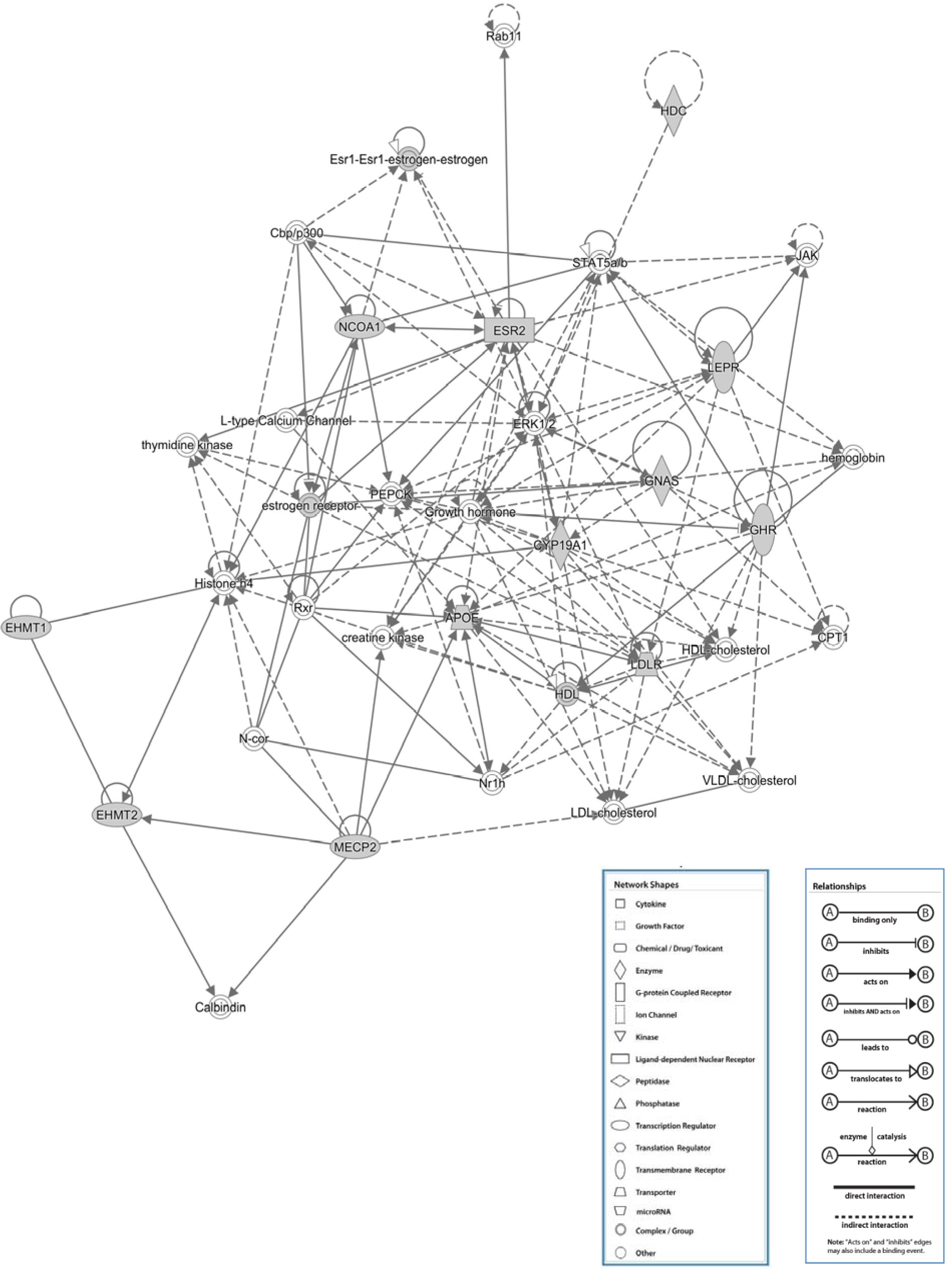
Pathways common to obesity and dementia. Ingenuity pathway analysis demonstrates that 12 of the 27 genes identified to be at the intersection of two often co-occurring conditions, obesity and dementia, map to one pathway.

### Finding pathways that underlie both life-extension drugs and dietary restriction that are conserved between *Mus musculus* and *Drosophila melanogaster.*

To determine whether a common network of genes underlies two lifespan extending interventions, that of dietary restriction or pharmacological intervention, we compared associated genomic data using GeneWeaver. Using data from studies of mouse or fly that examined gene expression in response to caloric restriction, the intersection of the two sets—one from each species—was taken using the “Boolean Algebra” tool in GeneWeaver. This intersection produced 45 cross-species homologs, representing the evolutionarily conserved targets of caloric restriction related to aging. Several drugs have been shown to extend lifespan in both species (see DrugAge database [25]) as representative drugs we selected Sirolimus-rapamycin, in mouse [26, 27], and 3,5,4'-trihydroxystilbene resveratrol, in fly [28] and using data from the CTD, a database of manually curated information about chemical gene/protein interactions, we sought to create a set of representative life extending drug target genes, one set of genes was created for each drug. Using data from each drug the intersection of the two sets—one from each species—was taken using the “Boolean Algebra” tool in GeneWeaver. To determine whether the 45 caloric restriction genes and these two drugs share a common molecular mechanism of action, the set of 45 homologs was overlapped with the data set produced from the two-way intersection of “life-extending drugs.” HSPA1A, an HSP70 complex member, was the one gene product common to all pathways. This is consistent with previous aging studies suggesting that the abundance of HSP70 complex members decreases with aging [7, 29], and identifying SNPs that are linked to HSP70 genes and are associated with longevity [30, 31].

This approach identified a gene product, HSPA1A, common to dietary restriction and two known drugs that influence life span. To find out whether there are additional drugs/chemicals that could also function to target gene products that underlie the beneficial effects of dietary restriction we used the GeneWeaver “View Similar GeneSets” tool to identify gene sets that are associated with other drugs that overlap with our set of 45 aging gene homologs. The top ten compounds interacting with any of the 45 genes of interest are shown in Table 2. These compounds need to be investigated for their direction of effect, as they may either shorten or extend lifespan, depending upon the direction of gene expression change they induce, and depending on the results of the corresponding diet-restricted study.

**Table 2.**
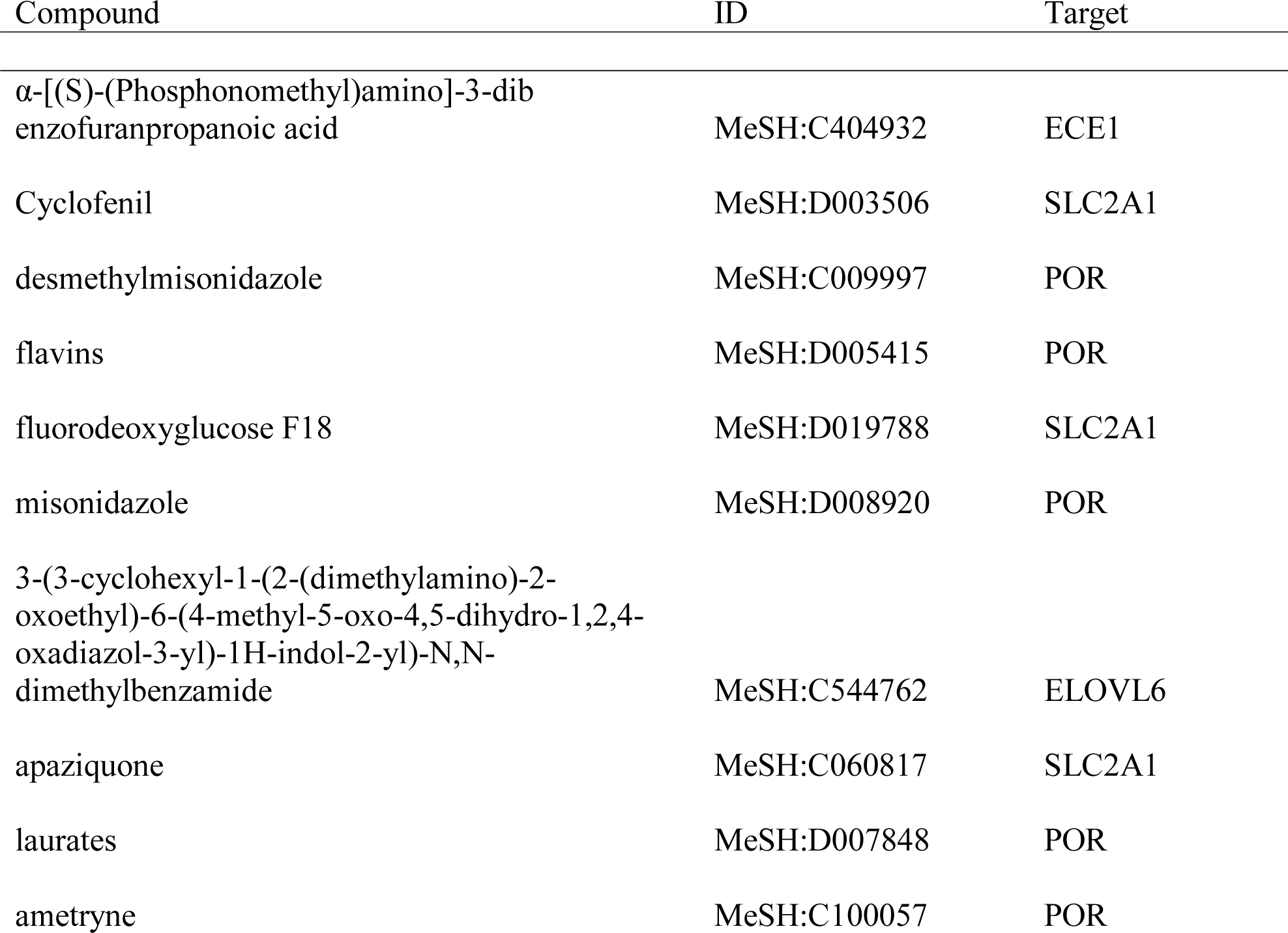
The top ten compounds known to interact with any of the 45 genes found at the intersection of cross-species dietary restriction aging studies.

### Cross-species translation of age-related processes

To identify convergent evidence across species for genes involved in aging, we integrated data from a total of 73 aging-associated gene sets (S2 Table), derived from 31 publications across 6 species (yeast, worm, fly, rat, mouse, human), and from three web resources (GeneNetwork, GenAge [32], and GWAS Catalog (https://www.ebi.ac.uk/gwas/). Using the “GeneSet Graph tool” in GeneWeaver, we identified *Cd63* as the most highly connected gene (i.e. it was present in the largest number of sets of genes) (Fig 3). *Cd63* was present in 12 gene sets from seven publications across four species (fly, rat, mouse, and human; Table 3). To validate *Cd63* as an aging gene, we knocked down the *C. elegans* ortholog, *tsp-7*, by feeding RNAi and observed a 5.6% extension of mean lifespan (18±4, n = 312 for *tsp-7(RNAi)* vs. 19± 5 days, n = 317 for *empty vector(RNAi);* p=0.00960 by the Wilcoxon Rank Sum test) (Fig 4, S3 Table).

**Fig 3.**
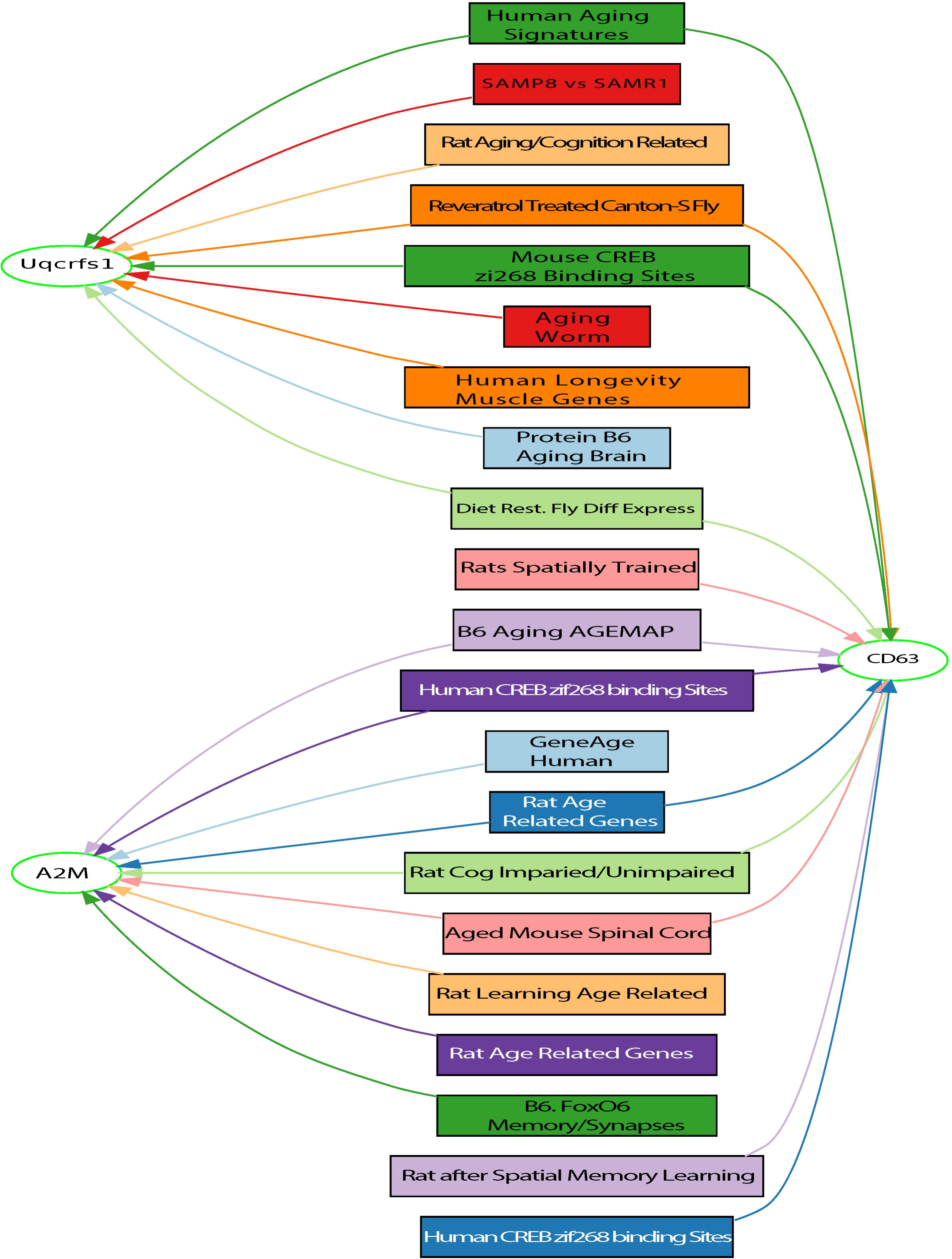
GeneSet graph of the most highly connected genes from 73 gene sets from six different species. The GeneSet Graph Tool presents a partitioned display of genes and GeneSets. Genes are represented by elliptical nodes, and GeneSets are represented by boxes. The least-connected genes are displayed on the left, followed by the GeneSets, then the more-connected genes in increasing order to the right. Genes and GeneSets are connected by colored lines to show what genes are in which GeneSets.

**Table 3.** Twelve of the 73 gene sets that contain *Cd63*.

| GeneSet Number | Species | GeneSet Name | Publication |
| --- | --- | --- | --- |
| 213295 | Human | Aging Signatures | [7] |
| 213272 | Fly | Resveratrol Canton-S | [42] |
| 216489 | Mouse | CREB zif268 binding sites | [43] |
| 216491 | Human | CREB zif268 binding sites | [43] |
| 216492 | Rat | CREB zif268 binding sites | [43] |
| 218922 | Rat | Aged Spatially Trained | [44] |
| 215692 | Mouse | Aged Spinal Cord | [45] |
| 213041 | Mouse | B6 Aging AGEMAP | [45] |
| 213271 | Fly | Diet-restricted Diff Expression | [42] |
| 137847 | Rat | Cognitive Impaired vs Unimpaired | [46] |
| 218921 | Rat | Age-related Genes | [47] |
| 218978 | Rat | Spatially Trained | [44] |

**Fig 4.**
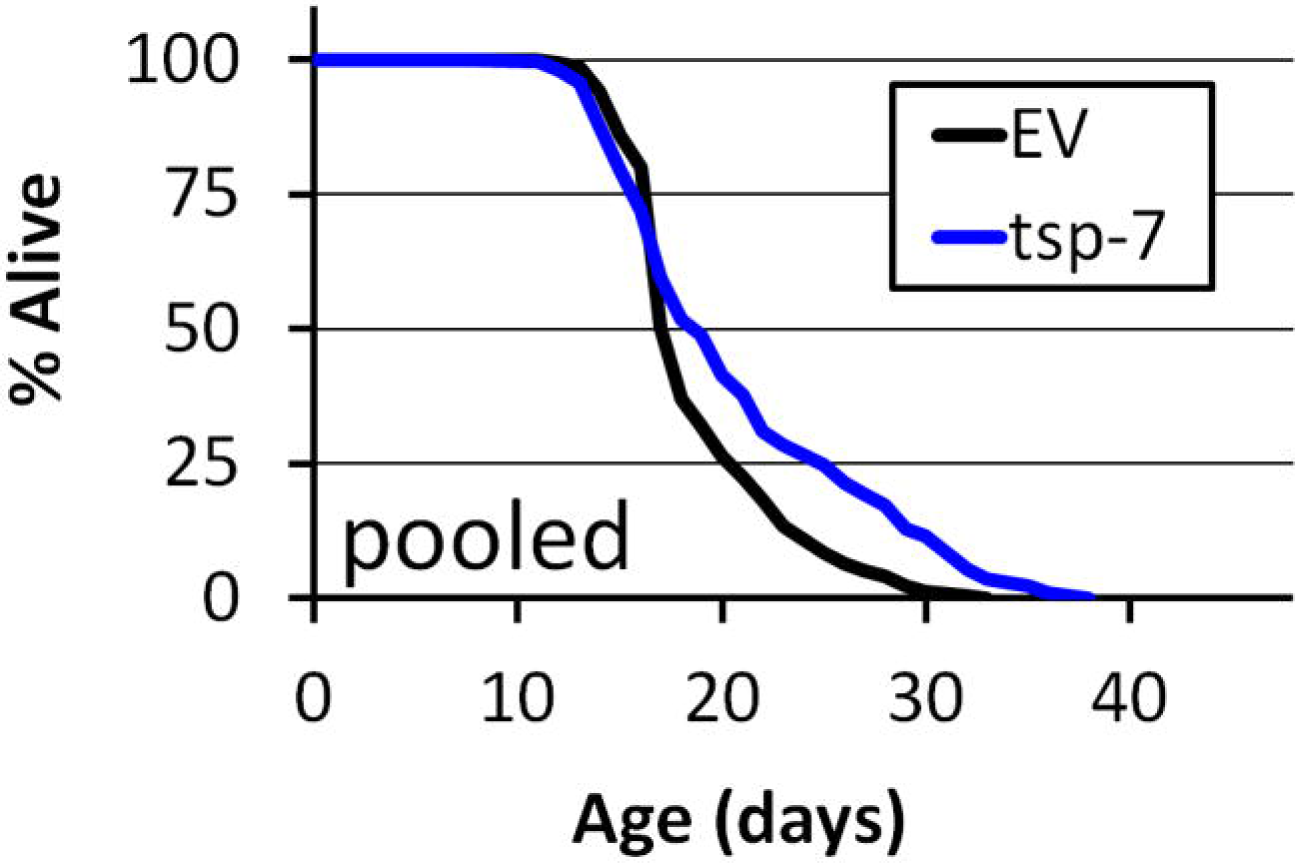
RNAi knockdown of *tsp-7* increases worm lifespan. Survival curves *C. elegans* fed either *empty vector (EV) RNAi* (black, n = 317) or *tsp-7(RNAi)* (blue, n = 312).

Further inspection of the same GeneSet graph (Fig 3) showed that ubiquinol-cytochrome c reductase, Rieske iron-sulfur polypeptide 1 *(Uqcrfsl)* is present in nine sets. Rieske iron-sulfur proteins have been linked to aging in yeast [33] and worms [34, 35]. Furthermore, *Uqcrfs1* is involved in mitochondrial function [36], and a decline in activity and quality of mitochondria is associated with age-related diseases [37]. The third gene on that same graph is alpha-2-macroglobulin (A2M). A2M has a well-characterized role in Alzheimer’s disease and aging in multiple species [38-40]. Together, these results demonstrate the utility of integrative functional genomics to identify aging-related genes using integrative gene set analysis across multiple species.

### Evaluation of CD63 in human GWAS studies

To determine if *Cd63* has been significantly associated with aging phenotypes in the recent large GWAS studies by the UK Biobank, we downloaded data from LD Hub and looked for association of SNP in or *near Cd63* with age related phenotypes (age of father’s death, age of mothers death etc). The UK Biobank aging dataset contained five SNPs which occur in CD63. Three (rs2231462, rs142309837, rs3138132) of these SNPs are 5’ UTR variants, one (rs2231464) is an intron variant, and another (rs35746357) is a noncoding transcript exon variant. However, none of these were significant for any of the three aging phenotypes we examined. We also examined SNPs immediately downstream and upstream of CD63. There were nine downstream and 11 upstream variants, none of these were significant. The most significant upstream variant, rs144565701 which is roughly 4KB from *Cd63*, was for the “mother’s age at death” phenotype (rs144565701, p=0.034, q=0.85).

## Discusssion

The growing number of studies and data in many fields, including ageing, require the development of integrative and computational approaches to analyze the data. Using GeneWeaver’s database and analysis tools to address questions in aging research we were able to identify genes common to cellular senescence and functional cognitive decline; to examine gene products at the intersection between obesity and dementia, to identify several druggable targets for longevity, and to identify and validate a cross-species age-related gene from convergent evidence. Our identification of the role for CD63 in aging would not have been made without this use of this large genomic analysis tool. CD63 in *C.elegans* is member of the tertaspanin family of proteins. Recently it was shown that tsp-3 in *C.elegans* greatly extend (>20% extension) lifespan as well. Tetraspanis are transmembrane scaffolding proteins involved in motility, cell adhesion, proliferation and activation.

As more aging-related functional genomic data is generated and made public by scientists all over the world, integrative functional genomics strategies will allow efficient integration of each new study into the growing pool of meta-data and rapid analysis that leverages the diversity of data produced across species and technical disciplines. Here we used several analytical approaches available in GeneWeaver, while other applications of GeneWeaver are available beyond these examples, including prioritization of QTL positional candidates, integration of GWAS data with expression data, and identification of animal models of aging phenotypes based on their underlying biology. These and other applications can be readily executed in GeneWeaver by users and have been summarized in a recent publication [41]. The GeneWeaver database continues to grow in both the number and the variety of gene-sets it contains. Investigators in the aging research community are encouraged to submit their own studies to this system. The power and scope of integrative tools like GeneWeaver expand with each new user and dataset. GeneWeaver tools and data resources are in continued development that will allow for integration of heterogeneous pathway-centric data, integrating at the level of pathway rather than at the level of genes, across experiments and species. Together, these advances in aging data resource aggregation and analytics will enable the aging research community to readily identify convergent molecular evidence for novel mechanisms of aging, healthspan and lifespan.

## Acknowledgements

We thank Stephen Sampson for his feedback on this manuscript. We appreciate the contributions of previous Chesler lab members in the curation of the aging literature in GeneWeaver specifically; Kathryn Toal, Courtney M. Vaughn, Laura C. Anderson. This work was supported by NIH R01 AA018776 and Nathan Shock Center of Excellence on the Biology of Aging NIH AG038070.

---

**S1 Table- The sets of genes derived from eight published studies on differentially expressed genes associated with either functional decline or aging-related cognition and memory phenotypes.**

**S2 Table- The 73 aging-associated gene sets used to identify Cd63 and associated publications.**

**S3 Table- Raw data from the *C. elegans* fed either *empty vector (EV) or tsp-7(RNAi)***

